# Analysis of gene expression in *Aedes aegypti* suggests changes in early genetic control of mosquito development

**DOI:** 10.1101/2024.12.02.625715

**Authors:** Renata Coutinho-dos-Santos, Daniele G. Santos, Lupis Ribeiro, Jonathan J. Mucherino-Muñoz, Marcelle Uhl, Carlos Logullo, A Mendonça-Amarante, M Fantappie, Rodrigo Nunes-da-Fonseca

## Abstract

*Aedes aegypti,* a critical vector for tropical diseases, poses significant challenges for studying its embryogenesis due to difficulties in removing its rigid chorion and achieving effective fixation for in situ hybridization. Here, we present novel methodologies for fixation, dechorionation, DAPI staining, and in situ hybridization, enabling the detailed analysis of gene expression throughout *Ae. aegypti* embryogenesis. By synchronizing eggs at various developmental stages (0–72 h), we localized the transcripts of the gap gene *mille-pattes (mlpt),* the dorsoventral gene cactus *(cact),* and the pioneer transcription factor (pTF) *zelda (zld).* In situ hybridization and RT-qPCR analyses revealed that *mlpt and cact* are maternally expresse*d, while zld* expression begins zygotically during cellularization and later becomes prominent in neuroblasts. Analysis of previously published transcriptomes suggests that three other pTFs, *CLAMP, grainyhead* and *GAF,* are also maternally expressed and may function as pioneer transcription factors during *Ae. aegypti* embryogenesis. These findings suggest that the transcription factors responsible for genome activation in mosquitoes differ from those in fruit flies, highlighting significant divergence in the genetic regulation of early Dipteran embryogenesis.

## INTRODUCTION

The yellow fever mosquito, *Aedes aegypti*, is a primary vector for several arboviruses, including Dengue, Zika, and Chikungunya, particularly in tropical and subtropical regions (Kraemer et al. 2019). In recent years, efforts by various research groups have successfully established *Ae. aegypti* as an emerging insect model organism, leveraging technologies such as Cas9/CRISPR-mediated genome editing alongside other technical advancements (Matthews and Vosshall 2020)

While extensive knowledge exists about the larval, pupal, and adult stages of *Ae. aegypti*, relatively few studies have focused on its embryogenesis (Christophers 1960, Vital et al. 2010, Farnesi et al. 2019, da Silva et al. 2021, Mundim-Pombo et al. 2021, Reid et al. 2024, Schember et al. 2024). Most cellular and molecular insights into insect embryogenesis have been derived from studies of the fruit fly, *Drosophila melanogaster*. However, the common ancestor of mosquitoes (Culicomorpha) and fruit flies (Cyclorrhapha) lived approximately 220 million years ago, underscoring the evolutionary divergence between these two groups (Yeates and Wiegmann 2017).

Although both *Drosophila melanogaster* and *Aedes aegypti* embryos are of the long-germ type - where all body segments are specified simultaneously along the entire egg length (Davis and Patel 2002) - significant differences exist in their embryogenesis. First, while mesoderm invagination occurs simultaneously along the entire anteroposterior axis in *D. melanogaster*, classical histological studies suggest that mesoderm internalization in *Ae. aegypti* does not take place by invagination (Christophers 1960). Second, *Ae. aegypti* embryos possess two distinct extra-embryonic membranes: the amnion and the serosa. The serosa surrounds the embryo during late developmental stages, whereas in *D. melanogaster*, the amnioserosa is a transient structure that degenerates before larval hatching (Rafiqi et al. 2012). Third, recent evolutionary analyses of anteriorly and posteriorly localized mRNAs involved in embryonic polarity in Diptera revealed that the anterior determinant in *Ae. aegypti* is the C2H2 zinc-finger gene *cucoid* (Yoon et al. 2019). In contrast, this role in *D. melanogaster* is performed by *bicoid*, a duplicated Hox3 ortholog found exclusively in cyclorrhaphan flies (Stauber et al. 2002).

Investigating the spatial analysis of gene expression during the early embryogenesis of *Aedes aegypti* has been notably challenging, primarily due to difficulties in removing the rigid chorion without damaging the embryo and its extraembryonic membranes (Christophers 1960, Vital et al. 2010, Farnesi et al. 2019, da Silva et al. 2021, Mundim-Pombo et al. 2021, Reid et al. 2024, Schember et al. 2024). Consequently, establishing reliable methods for fixation and in situ hybridization of *Ae. aegypti* embryos that encompass all stages of embryogenesis remains a significant hurdle.

Here, we present a new method for fixing *Ae. aegypti* embryos and studying spatial gene expression using *in situ* hybridization, with a particular focus on the earliest developmental stages. Utilizing this approach, we successfully localized the expression of the gap gene *mille-pattes (mlpt)*, the dorsoventral patterning gene *cactus (cact)*, and the pioneer transcription factor (pTF) *zelda (zld)*.

Comparison of *in situ* hybridization results with real-time quantitative PCR (RT-qPCR) data and previously published transcriptomes revealed that the pTF *zld* is not maternally deposited, suggesting that genome activation in *Ae. aegypti* likely involves other pTFs. Further analysis of transcriptomes identified orthologs of *CLAMP, trl (GAGA), and grainyhead (grh)* as both maternally provided and zygotically transcribed, indicating that they might play key roles as pioneer transcription factors in this mosquito.

## MATERIALS AND METHODS

### Maintenance of *Aedes aegypti* Colony

The *Aedes aegypti* Rockefeller strain was maintained under controlled conditions: 80% humidity, 28°C, and a 12-hour light/dark cycle in an incubator (411FPD 335, Ethiktechnology) at the Hatisaburo Masuda Integrated Biochemistry Laboratory, part of the Institute of Biodiversity and Sustainability (NUPEM) at the Federal University of Rio de Janeiro. Adults were housed in plastic cages covered with fine tulle mesh and provided ad libitum access to a 10% sucrose solution. Larvae were kept in plastic trays filled with water and fed on rat food, with the trays also covered by tulle mesh.

Female mosquitoes were blood-fed using immobilized mice positioned on top of the cages, with their abdominal regions facing downward, for 20 minutes. All procedures involving vertebrates adhered to ethical standards set by the Animal Use Ethics Committee, with mosquito feeding approved under protocol number MAC002.

### Forced Induction of Female Oviposition

Oviposition was conducted 72 hours post-blood-feeding, either naturally or through forced induction. For forced oviposition, females were transferred to Petri dishes (90 x 15 mm) lined with filter paper moistened with distilled water. The dishes were covered with aluminum foil to prevent light exposure and placed in an incubator (28°C, 80% humidity). Within 15 minutes, females began laying eggs, marking the zero-hour starting point. Eggs synchronized at specific stages (0–1h, 1–2h, 2–3h, etc.) were collected, transferred to 1.5 mL microtubes, fixed, and stored at -20°C to halt embryonic development.

For later-stage eggs (e.g., 15–16h, 19–20h, 23–24h, etc.), natural oviposition was performed within the cages. Wet filter paper was introduced, allowing females to lay eggs naturally for 75 minutes. Following egg collection, the females were discarded.

### Fixation of *Ae. aegypti* Eggs

Eggs at all stages were transferred from Petri dishes to 1.5 mL microtubes using a fine paintbrush and covered with distilled water. The microtubes were submerged in a boiling water bath for 90 seconds, then immediately transferred to an ice bath for another 90 seconds to induce thermal shock. Eggs were then placed in glass screw-cap tubes containing a 1:1 mixture of 4% paraformaldehyde (PFA) and heptane (Vetec). The tubes were shaken at maximum speed (Kline NT 151 shaker, Novatecnica) for one hour. Following fixation, the PFA was replaced with methanol, and the samples were agitated again to ensure thorough mixing. Eggs were washed twice in methanol and stored at -20°C for *in situ* hybridization and nuclear labeling with DAPI (4′,6-Diamidino-2-fenilindol, 2-(4-Amidinofenil)-6-indolecarbamidin-Sigma-Aldrich).

### Chorion Removal of *Ae. aegypti* Eggs

Fixed eggs were gradually transferred to phosphate-buffered saline with Tween 20 (PBST) to prevent osmotic shock. Eggs were placed onto filter paper (2 x 6 cm) lightly moistened with PBST and carefully adhered to double-sided tape affixed to a Petri dish (90 x 15 cm). Once the eggs were deposited, a 1x PBS (Phosphate Buffer Solution) solution was added to cover them, ensuring their integrity was maintained. Under a stereoscope (Olympus SZ2-LGCL), chorions were removed manually using Dumont No. 4 forceps by gently tapping and peeling away the rigid outer layer. The dechorionated eggs were transferred into 1.5 mL microtubes containing PBST using a 200 µL pipette with a trimmed tip to prevent damage. Following this, the eggs were gradually returned to methanol and stored at -20°C for subsequent *in situ* hybridization and DAPI staining.

### RNA Isolation, Purification, and cDNA Synthesis

RNA isolation was carried out using Trizol® reagent (Ambion by Life Technologies), following the manufacturer’s protocol. Total RNA was extracted from *Ae. aegypti* eggs, using a quantity equivalent to 0.1 mL of eggs in a 1.5 mL microtube, to which 500 µL of Trizol® reagent was added. Subsequently, 1 µL of the isolated RNA was used to assess concentration and purity using a NanoDrop 2000c spectrophotometer (Thermo Scientific). The purity of the RNA was evaluated based on the absorbance ratios at 260 nm / 280 nm and 260 nm / 230 nm. Complementary DNA (cDNA) synthesis was performed using the High-Capacity cDNA Reverse Transcription Kit (Applied Biosystems), following the manufacturer’s instructions. The samples were then placed in a Veriti 96-Well Thermal Cycler (Applied Biosystems) for cDNA synthesis.

### Identification of Early Expressed Orthologs in the *Ae. aegypti* Genome

Ortholog identification in the *A. aegypti* genome was performed using OrthoDB (v11) (Kuznetsov et al. 2023). Verification was achieved through BLASTP (Altschul et al. 1990) and reciprocal BLAST searches (http://blast.ncbi.nlm.nih.gov/). Orthologs of pioneer transcription factors (pTFs) identified include *zelda* (zld-173608at7147, reported by Ribeiro et al. 2017), *Clamp* (125906at7147), *grainyhead* (*grh-10646at7147*), *Trithorax-like* (*Trl-GAGA-203803at7147*), and *Odd paired* (*Opa-169368at7147*). The dorsoventral gene *cactus* (*cact-195346at7147*) was also identified, alongside the gap gene *mille-pattes* (*mlpt*), previously reported in *Ae. aegypti* by Tobias-Santos et al. 2019.

### Primer Design, PCR Amplification, and Probe Synthesis

Primers for in situ hybridization were designed using Primer3 (http://frodo.wi.mit.edu/) and synthesized by Invitrogen. Product sizes and sequences are detailed in Supplementary Table 1. Forward and reverse primers included specific 8-nucleotide adaptor sequences at their 5’ ends to allow T7 adaptor addition. Template amplification for in situ hybridization probes involved two successive PCR reactions. The first PCR used embryonic cDNA pools as templates. The second PCR utilized an aliquot of the first PCR product as a template, employing the same forward primer along with a T7 universal primer (3’-agggatcctaatacgactcactatagggcccggggc). Eight microliters (∼100-200 ng) of the second PCR product served as a template for the synthesis of antisense RNA probes. This synthesis was carried out using T7 RNA polymerase and Dig RNA Labeling Mix (Roche), yielding labeled probes for in situ hybridization experiments.

### Whole-Mount *In Situ* Hybridization

Dechorionated embryos were gradually rehydrated from methanol to 1× PBST (PBS with 0.1% Tween 20), followed by a 5-minute wash on a K45-3220 Blood Mixer shaker (Kasvi). The embryos were fixed in 4% paraformaldehyde (PFA, w/v) for 10 minutes under constant agitation. Post-fixation, three 5-minute washes with 1× PBST were conducted before redistributing the embryos into pre-labeled 1.5 mL microtubes. The embryos were transferred stepwise into a hybridization solution (Hyb II) supplemented with salmon sperm DNA, with agitation for 5 minutes at each transition (50% 1× PBST/50% Hyb II, followed by 25% 1× PBST/75% Hyb II). Two washes in 100% Hyb II were performed, each lasting 5 minutes, before pre-hybridization at 55°C for 1 hour in a dry bath. Subsequently, 500 µL of the solution was removed, and 2 µL of previously synthesized probes were added for hybridization at 55°C overnight. Post-hybridization, embryos were washed four times with pre-warmed Hyb II for 30 minutes at 55°C. Gradual rehydration to 1× PBST (2:1, then 1:2) followed, with 15 minutes for each step. After a wash with 1× PBST, the embryos underwent blocking with 1:5-diluted Western Blocking Solution (Roche) for 1 hour at room temperature under agitation. The blocking solution was replaced with a fresh solution containing Anti-digoxigenin-AP antibody (Roche) at a 1:5000 dilution, and the embryos were incubated overnight at 4°C. After returning to room temperature, the embryos were washed twice with 1× PBST, followed by three additional 15-minute washes. An alkaline phosphatase (AP) buffer (Tris pH 9.5, MgClL, NaCl, Tween 20) was used for one 5-minute wash, followed by two more washes with the same buffer. Color development was initiated by immersing embryos in BM-Purple Solution (Roche) within 24-well plates, monitored for 30 minutes to 48 hours. To stop the reaction, embryos were washed twice with 1× PBST and stained with DAPI (1 µg/mL) for 10 minutes. The stained embryos underwent two final washes with 1× PBST before being analyzed under a Leica M205 FA Stereo microscope equipped with fluorescence.

### Quantitative Gene Expression Analysis Using RT-qPCR

RNA was isolated from *Ae. aegypti* embryos at specified developmental stages (0-1h, 4-5h, 5-6h, 23-24h, 47-48h, and 71-72h) using Trizol® reagent (Ambion by Life Technologies), following the manufacturer’s instructions. RNA purity was assessed using absorbance ratios at 260/280 nm via a NanoDrop 2000c spectrophotometer (Thermo Scientific). Reverse transcription was performed on 2 µg of total RNA using the High-Capacity cDNA Reverse Transcriptase Kit (Applied Biosystems). RT-qPCR primers for *zld*, *mlpt*, and *cact* were designed with Primer3 (http://frodo.wi.mit.edu/), with sequences detailed in Supplementary Table 1. SYBR Green assays (qPCRBIO SyGreen^®^ Mix) were run on the QuantStudio 3 platform (Applied Biosystems), following the manufacturer’s protocol. The results, which were recorded as Ct (threshold cycle) values, were converted into relative expression values using the methods “ΔCt” (Schmittgen and Livak, 2008) for ovary and carcasses and “ΔΔCt” (Livak and Schmittgen, 2001) for embryos. Both analyses were normalized to rp49 transcript levels (Gentile et al. 2005, Amarante et al. 2022).

### Statistical analysis of gene expression

Statistical analysis was performed using the mean of three independent experiments. The Kruskal-Wallis test followed by Tukeýs multiple comparisons test was used for statistical analysis of ovary and carcass samples. For embryonic samples, Dunnett’s multiple comparisons post-test was used. Differences were considered significant whenever p ≤ 0.05. GraphPad Prism 6 software (v. 6.01; GraphPad software Inc., San Diego, CA, USA) was used for statistical analysis.

### Bioinformatic Analysis of Published Transcriptomes

Expression profile of twenty public RNAseq libraries of *Ae. aegypti* were recovered from NCBI/SRA (https://www.ncbi.nlm.nih.gov/sra). Libraries representing the initial stages of embryonic development of *Ae. aegypti* were downloaded. Four developmental points of whole embryos were recovered in quadruplicates: one hour of development (ERR9736484, ERR9736485, ERR9741330 and ERR9741331), three hours of development (ERR9736486, ERR9736487, ERR9741332 and ERR9741333), five hours of development (ERR9736488, ERR9736489, ERR9741334 and ERR9741335) and seven hours of development (ERR9736490, ERR9736491, ERR9741336 and ERR9741337). Transcriptomes of 0-1 hour egg anterior halves (SRR8729853 and SRR8729854) and posterior halves (SRR8729851 and SRR8729852) were also used for transcriptome analysis. The binary files of the libraries were transformed to FastaQ format using SRAToolKit(v3.1.1) (Team 2021). Trimmomatic (v0.39) (Bolger et al. 2014) was used for trimming of adapters residues based on each sequencing technology of each library and quality filtering of the reads by Phed33. The transcriptome of the most recent version (AaegL5.0) of the *Ae. aegypti* reference genome (GCF_002204515.2) strain LVP_AGWG was used for the construction of an index using the Salmon-Index function of Salmon (v1.10.1) (Patro et al. 2017). To quantify the expression level in the libraries in transcripts per million (TPM), the reads of the libraries were mapped against the index/transcriptome using the Salmon-Quant function of Salmon (Patro et al. 2017). The TPM values per gene and development time were obtained and plotted. *zld*, *cactus* and *mille-pattes*, and other four putative pioneer transcription factors *Chromatin-linked adaptor for MSL proteins (Clamp), Grainy head (grh), Trithorax-like transcription factor GAGA (Trl-GAGA) and Odd paired (opa)* were used for the analysis.

## Results and Discussion

### Visualization of *Ae. aegypti* Embryonic Development via New Fixation Method

Fixing early *Aedes aegypti* embryos for visualization has been historically challenging due to the rigidity of the chorion, which often leads to degradation of embryos during processing. This limitation has restricted the ability to perform techniques such as *in situ* hybridization, particularly during early developmental stages. Previously established methods (Rezende et al. 2008, Farnesi et al. 2009, Clemons et al. 2010, Vargas et al. 2014, Farnesi et al. 2019, Mundim-Pombo et al. 2021) have used various chemicals to degrade the chorion, but these often result in compromised sample integrity.

In this study, we introduce a novel fixation method that addresses these limitations, allowing for precise nuclear staining with DAPI and successful *in situ* hybridization. This method provides high-quality visualization of *Aedes aegypti* embryogenesis, from initial cleavages to the formation of larval morphology at 72 hours post-oviposition, incubated at 28°C (Figure 1).

**Figure 1:**
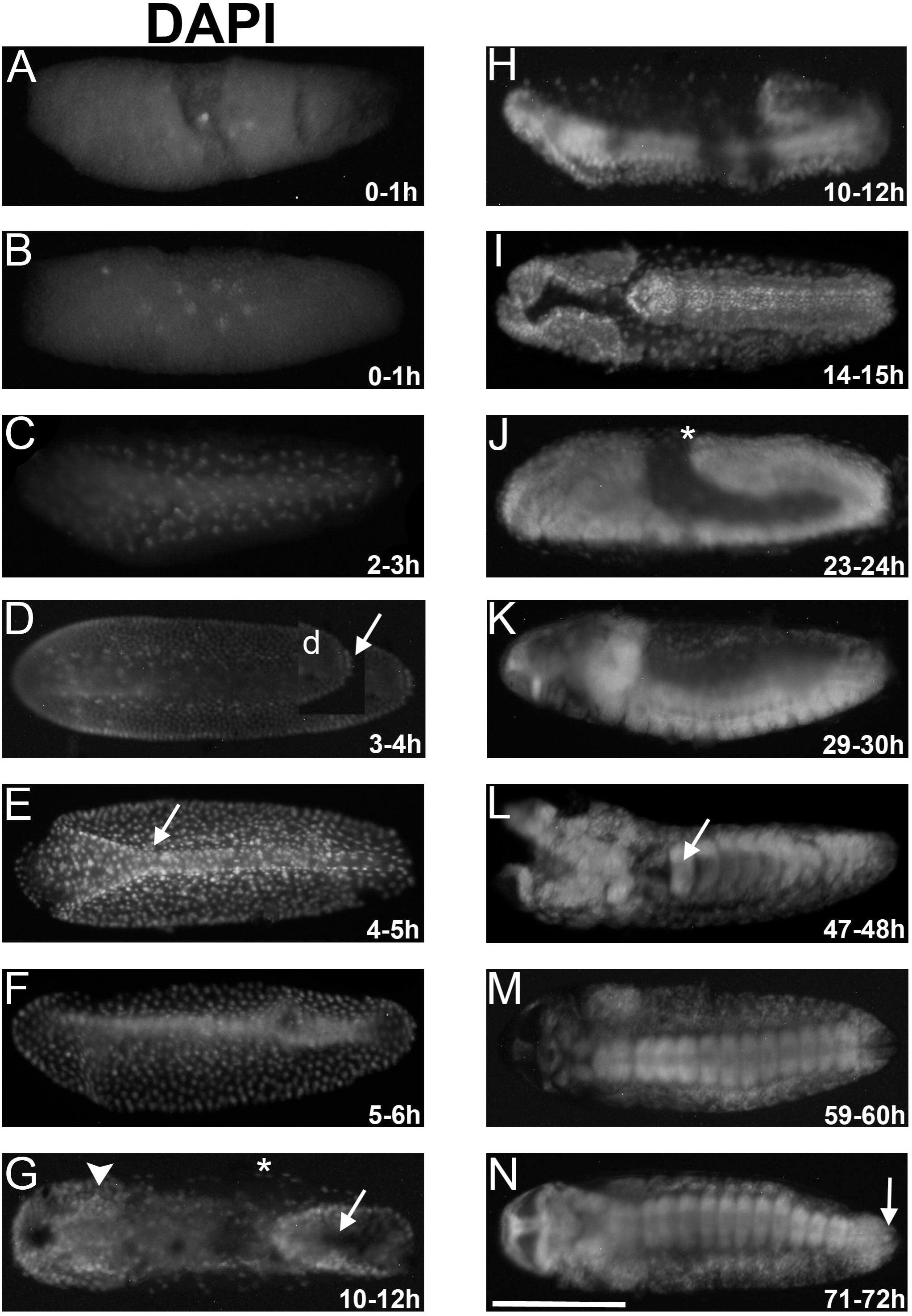
Embryonic development of *Aedes aegypti* using DAPI staining. Staining at 0-1h after egg laying (AEL) showing the first cleavages. About 5 nuclei (A) and 9 (B) can be observed. (C) The nuclei increase in number and migrate to the periphery at 2-3h AEL. (D) Lateral view. The inset indicates the posterior region where germ cells can be identified due to their unique rounded shape (arrow). (E) Ventral view. An embryo at 4-5 hours AEL with two distinct regions of mesoderm internalization, the anterior region is broader (arrow) than the posterior one. (F) Ventral view. At 5-6h after the invagination process is taking place, with more cells than seen in previous stages. (G) At 10-12 hours AEL, dorsal view. An embryo during germ band extension (arrow) showing a well-defined head (arrowhead), and the extra-embryonic serosal cells (asterisk). (H) At 10-12 hours AEL, lateral view of the same embryo from the previous image (G) during germ band extension process. (I) At a 14-15h AEL, dorsal view. The embryo is further extending the germ band. (J) At 23-24 hours AEL, lateral view. During germ band retraction stage, the serosal cells can be observed at the dorsal egg region (asterisk). (K) At 29-30 hours AEL, lateral view. An embryo starting the dorsal closure process. (L) At 47-48 hours AEL, ventral view. Nervous system is morphologically distinct at the ventral region of the embryo. (M) At 59-60 hours AEL, ventral view. An embryo whose head segments appear fully segmented, and the open stomodeum is clear. The thoracic and abdominal segments remain similar in width. (N) At 71-72 hours AEL, ventral view. A clear distinction between thoracic and abdominal segments is evident and the respiratory siphon (arrow) is observed at the posterior region.

Embryonic development begins with internal meta-synchronous nuclear cleavages (0 to 2 hours after egg laying, AEL), during which the nuclei migrate to the egg periphery (Figure 1 A,B). Between two to three hours after egg laying (AEL) these nuclei start to be seen at the periphery (Figure 1 C) and between three to four hours AEL, the number of nuclei has drastically increased and a distinct cell population corresponding to the pole cells can be observed at the posterior most region of the egg (Figure 1 D, d-arrow). A recent study showed via antibody staining’s that these posterior cells express the germ cell specific factor *vasa* (Zhang et al. 2023).

Between four to five hours AEL, gastrulation, e.g. mesoderm internalization takes place (Figure 1 E, -arrow). In contrast to *Drosophila melanogaster* where mesoderm invagination is uniform and occurs along the whole antero-posterior axis, other processes such as individual cell ingression have been reported in other insects. Three different mesodermal internalization types have been identified in insects and, in the beetle, *Tribolium castaneum* these three types can be recognized in a single embryo undergoing gastrulation (Roth 2004, Lemke et al. 2020). Analysis of DAPI staining of *Ae. aegypti* embryos suggest that two prospective mesodermal cell populations seem to exist, one at the anterior region with a wider open area, and a second region with a narrower opening located immediately posteriorly (Figure 1 E,F). Between 5 to 6 hours AEL, the mesoderm appears internalized, and a cephalic furrow is evident at the ventral side (Figure 1 F, arrowhead).

Shortly after, the embryo starts germ band elongation, which is evident between 11 to 12 hous AEL. During this process, the posterior embryonic region folds inwards like in *Drosophila melanogaster* and the germ cells are internalized and pushed (Figure 1 G,H). Between 14 to 15 hours AEL the embryo further elongates, as observed by the posterior progression of the posterior most zone (Figure 1 I). Between 23 and 24 hours AEL the embryo begins germ band retraction (Figure 1 J). By 29 and 30 hours the distinction between the head/gnathal and the thoracic segments becomes more evident (Figure 1 K). At 47 and 48 hours AEL the ventral nerve cord (arrow) is evident as well during late organogenesis, alongside the segmentation of the larvae (Figure 1 L). Between 59- and 60-hours AEL, the head segments appear fully segmented, the open stomodeum becomes evident, and the thoracic and abdominal segments remain similar in width (Figure 1 M). Shortly before hatching, at 71 to 72 hours AEL, the embryo acquires a “mosquito larva-like morphology,” characterized by a clear distinction between thoracic and abdominal segments and the emergence of the respiratory siphon (arrow) (Figure 1 N). This fixation protocol not only enables detailed visualization of Aedes aegypti embryogenesis from early to late stages but also provides a robust framework for comparative developmental studies. By facilitating precise analyses of developmental processes, it opens avenues for exploring evolutionary mechanisms and biological diversity within mosquitoes and related species.

### *In situ* hybridization pattern of *mlpt, cact*, and *zld* genes during *Ae. aegypti* embryogenesis

Since our fixation method enables the proper visualization of embryonic structures, we investigated the localized expression of key developmental markers during *Ae. aegypti* embryogenesis. Specifically, we examined the orthologs of the gap gene *mille-pattes (mlpt),* the dorsoventrally regulated gene *cactus (cact)* and the pioneer transcription factor *zelda (zld)*.

The *mlpt* gene encodes a polycistronic messenger-RNA (mRNA) with several small open reading frames (smORFs). It has been reported as a *gap* gene during beetle and bug embryogenesis (Savard et al. 2006, Ray et al. 2019, Tobias-Santos et al. 2019, Silva et al. 2024). In *D. melanogaster*, its ortholog *tarsaless* (*tal*) does not function as a gap gene but instead has a late role in trichome formation (Galindo et al. 2007).

Interestingly, distinct *mlpt* expression begin between 2 and 3 hours after egg laying (AEL), forming a broad domain in the middle of the egg (Figure 2 A,Á). By 3 to 4 hours AEL, this domain narrows, overlapping with the mesoderm and excluding the posterior germ cells (Figure 2 B,B’,C,Ć). From 4 to 5 hours AEL, the domain further narrows and becomes concentrated in the egg’s mid-region (Figure 2 D,D’). By 5 to 6 hours AEL, two stripes emerge in the mid-embryo, coinciding with posterior invagination (Figure 2 E,É,F,F’).

**Figure 2:**
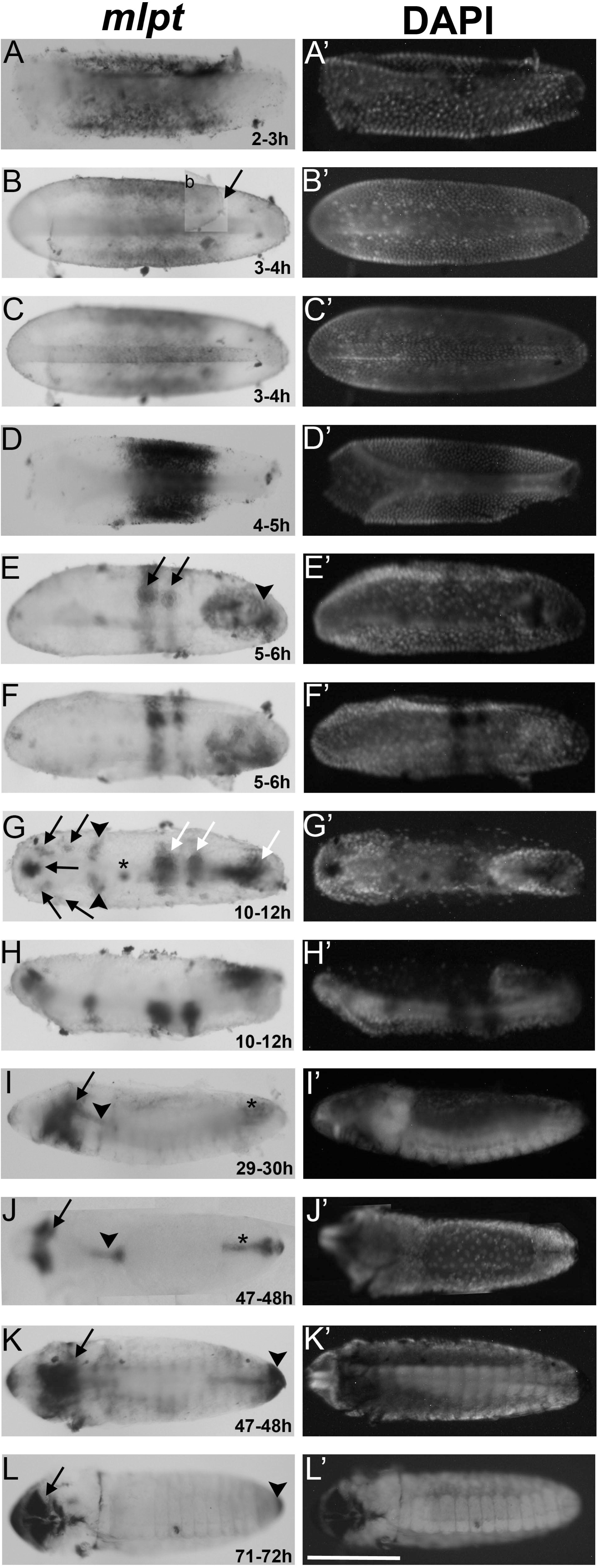
Expression pattern of the smORF-containing gene *mlpt* during *Aedes aegypti* embryonic development. (A, A’) Lateral view, early-stage embryo 2–3 hours AEL showing diffuse expression in the middle region. (B, B’) Ventral view, embryo at 3–4 hours AEL with more centralized staining in the middle region, extending towards the embryo’s periphery. A boxed inset highlights the posterior region where germ cells are evident (arrow) and do not express *mlpt*. (C, C’) Ventral view, embryo at 3–4 hours AEL showing that the mesodermal cells express *mlpt.* (D, D’) Ventral view, embryo at 4-5 hours AEL, the *mlpt* expression domain further narrows and becomes concentrated in the egg’s mid-region. (E, E’) Dorsal view of a 5–6-hour AEL embryo showing two distinct expression bands in the middle region (arrows) and strong posterior staining (arrowhead). (F, F’) Lateral view of the same 5–6-hour AEL embryo as in E, E’, highlighting the spatial expression patterns. (G, G’) By 11 and 12 hours AEL *mlpt* expression pattern shows a complex pattern with stripes in the middle-posterior region and localized expression in the anterior (stomodeum, head lobes, and gnathal segment). Discrete spots in the head (arrows), bands in the anterior/middle regions (arrowheads), a central spot (asterisk), bands in the posterior/middle region, and a prominent posterior domain (white arrows) are observed. (H, H’) Lateral view of the same 10–12-hour AEL embryo as in G, G’. (I, I’) Lateral view of a 29–30-hour AEL embryo showing expression in the head (arrow), anterior/middle head region (arrowhead), and posterior region (asterisk). (J, J’) Dorsal view of a 47–48-hour AEL embryo with pronounced staining in the prospective head (arrow), and in anterior/middle region (arrowhead), and posterior region (asterisk). particularly in the anterior and posterior regions of midgut undergoing invagination. (K, K’) Ventral view of the same 47–48-hour embryo as in J, J’, with strong expression in the anterior (head, arrow) and posterior (final segments, arrowhead) regions. (L, L’) Embryo at 71–72 hours with expression localized to the head (arrow) and the posterior-most region (arrowhead). Scale bar: 200 µm. Images captured using a Leica M205 stereomicroscope.

By 11 and 12 hours AEL *mlpt* expression pattern shows a complex pattern with stripes in the middle-posterior region and localized expression in the anterior (stomodeum, head lobes, and gnathal segment; Figure 2 G,G′,H,H′). At 29 to 30 hours AEL, expression persists in the anterior, middle/gnathal (arrowhead), and posterior (asterisk) regions (Figure 2 I,I′). Between 47 to 48 hours AEL (Figure 2 J,J’,K,K’), staining is observed in the head lobes and inside the embryo, particularly in the anterior and posterior regions of midgut undergoing invagination. This expression in tissues containing chitin such as the gut must be seen with caution, although experiments with sense probes and *cactus* and *zld* probes did not lead to similar staining patterns. Between 71 to 72 hours AEL expression cuticle formation is largely advanced particularly in the head and in the posterior, thus these staining’s might represent artifacts and must be seen with cautions (Figure 2 L,Ĺ).

Comparisons with other species suggest that the large gap domain observed during early development (2 to 5 hours AEL) is shared with several other species, such as the beetle *T. castaneum* and the hemiptera *R. prolixus* (Savard et al. 2006, Ray et al. 2019, Tobias-Santos et al. 2019). The stripe pattern in the middle of the embryo observed during gastrulation (5 to 6 hours AEL) might be similar to the seven blastodermal stripe pattern observed in *D. melanogaster* (Galindo et al. 2007), thus the early expression of *Ae. aegypti mlpt* display features of several species. Late expression at the head lobes, stomodeum and the posterior and anterior gut has also been observed in *R. prolixus* (Tobias-Santos et al. 2019).

Next, we examined the *in situ* hybridization pattern of the Ik-B ortholog *cact.* In *D. melanogaster cact* is maternally deposited and important to regulate the transport of NF-kB Dorsal between nuclei and cytoplasm during DV axis formation (Roth et al. 1991). In *D. melanogaster cact* is also expressed zygotically in the mesoderm, in the ventral most part of the *D. melanogaster* embryo under Dorsal and Twist control (Sandmann et al. 2007). In the short germ beetle *T. castaneum cact* is strictly zygotically expressed and participates in a zygotic feedback loop required for DV axis formation (Nunes da Fonseca et al. 2008). Thus, we analyzed *cact* expression during *Ae. aegypti* embryogenesis particularly focusing on the early stages of development. Distinct expression of *cact* is evident between embryos 2 to 3 hours AEL as a ventral stripe (Figure 3 A,Á), accompanied by faint maternal staining around the egg. This observation suggests that *cact* is also maternally supplied in *Ae. aegypti* as in *D. melanogaster*. Interestingly, *cact* is expressed along the whole anteroposterior axis, even though the anterior triangular domain is broader than the posterior one. Between 3 to 4 hours AEL, the domain remains at the ventral region (Figure 3 B,B’). Between 4 to 5 hours AEL the expression appears broader than the mesoderm, probably also extending to the non-neurogenic ectoderm (Figure 3 C,Ć,D,D’). Interestingly, at 5 to 6 hours AEL, the anterior triangular domain disappears as cells internalize, leaving a single ventral domain shortly before gastrulation (Figure 3 E,ÉF,F’,G,G’). By 20 to 21 hours AEL, during germ band elongation, *cact* is expressed below the ectoderm in the head (arrows), middle/anterior (asterisk), and posterior regions (arrowhead), as well as the serosa (white arrow; Figure 3H,H′). This expression pattern is largely different from *mlpt*, confirming the specificity of the methodology. Moreover, *cact* expression during *Ae. aegypti* embryogenesis suggests dorsoventral maternal and zygotic roles of this gene similarly to what has been reported for *D. melanogaster* and different from a strict zygotic role reported in wasps and beetles (Nunes da Fonseca et al. 2008, Buchta et al. 2013).

**Figure 3:**
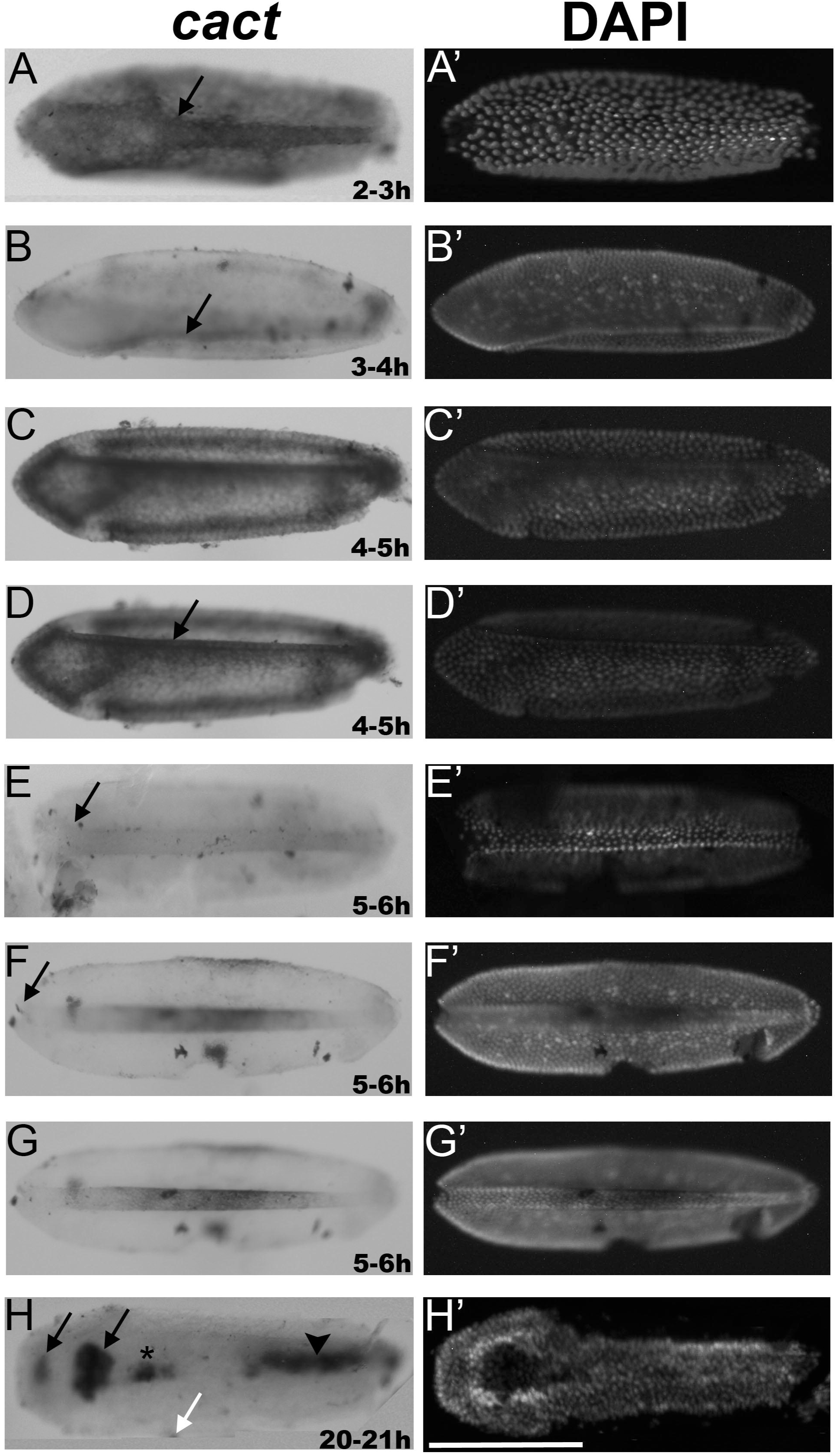
Expression pattern of the IκB inhibitor *cactus* during *Aedes aegypti* embryonic development. (A, A’) Ventral view, 2–3 hours AEL early-stage embryo showing staining in the prospective mesoderm (arrow) and diffuse expression throughout the embryo. (B, B’) Lateral view, an embryo at 3–4 hours AEL displaying staining at the ventral midline. (C,C’,D, D’) Ventral view, embryo at 4–5 hours AEL. Two different focal planes (C, D). A strong staining in the prospective mesoderm, which extends towards the periphery (neurogenic ectoderm) is evident. (E, E’, F, F’, G, G’) Ventral view, embryos at 5–6 hours AEL. *cact* expression concentrated in the middle region, the prospective mesoderm. (H, H’) Ventral view, embryo at 20–21 hours AEL showing prominent staining in the middle region, including the head (arrows), anterior/middle region (asterisk), and posterior internal region (mesoderm-arrowhead), particularly in areas associated with germ band formation. Additional diffuse staining is observed in the serosal layer (white arrow). Scale bar: 200 µm. Images captured using a Leica M205 stereomicroscope.

Lastly, the expression of the pioneer transcription factor *zelda (zld)* was analysed in *Ae. aegypti*. In *D. melanogaster zld* is maternally deposited and acts as an essential TF required for zygotic genome activation during MZT (Liang et al. 2008). In *D. melanogaster* Zld binds to a conserved heptamer motif enriched in regulatory regions of early expressed fruit fly genes (Harrison et al. 2011). In a few other insect species such as the beetle *Tribolium castaneum* (Ribeiro et al. 2017), the bee *Apis mellifera* (Pires et al. 2016), the wasp *Nasonia vitripennis* (Arsala and Lynch 2017) and the cockroach *Blatella germanica* (Ventos-Alfonso et al. 2019) *zld* is maternally deposited and early zygotically expressed, suggesting a conserved role in MZT during insect evolution. Thus, we expected that *zld* would be maternally expressed during oogenesis and embryogenesis during *Ae. aegypti* embryogenesis.

Remarkably, distinct *zld* expression is not detectable until 4 to 5 hours AEL, when cellularization begins. At this time point *zld* mRNA is broadly detected in the embryo, including the mesoderm (Figure 4 A,Á). During germ band elongation (14–15 hours AEL), strong expression appears in the head (arrow) and germ band (Figure 4 B, B’). By 20 to 21 hours AEL *zld* is observed in the anterior and posterior regions, respectively on the head (arrows) and germ band (arrowhead) and a staining on the serosal tissue (asterisk) (dorsal view in Figure 4 C,Ć and lateral view in Figure 4 D,D). Lastly, dorsal (Figure 4 E, E’) and lateral views (Figure 4 F, F’) of a 23-24 hours AEL embryo showed expression in the head (arrows) and along the embryo, specifically in the neuroblasts (arrowheads) in a parallel alignment (Figure 4 E, E’). Comparative analyses reveal conserved *zld* expression in neuroblasts across species, with additional posterior expression in short-germ species such as *B. germanica* and *T. castaneum* (Ribeiro et al. 2017, Reichardt et al. 2018, Ventos-Alfonso et al. 2019). The lack of maternal *zld* expression in *Ae. aegypti* observed by *in situ* hybridization prompted further investigations using alternative methods.

**Figure 4:**
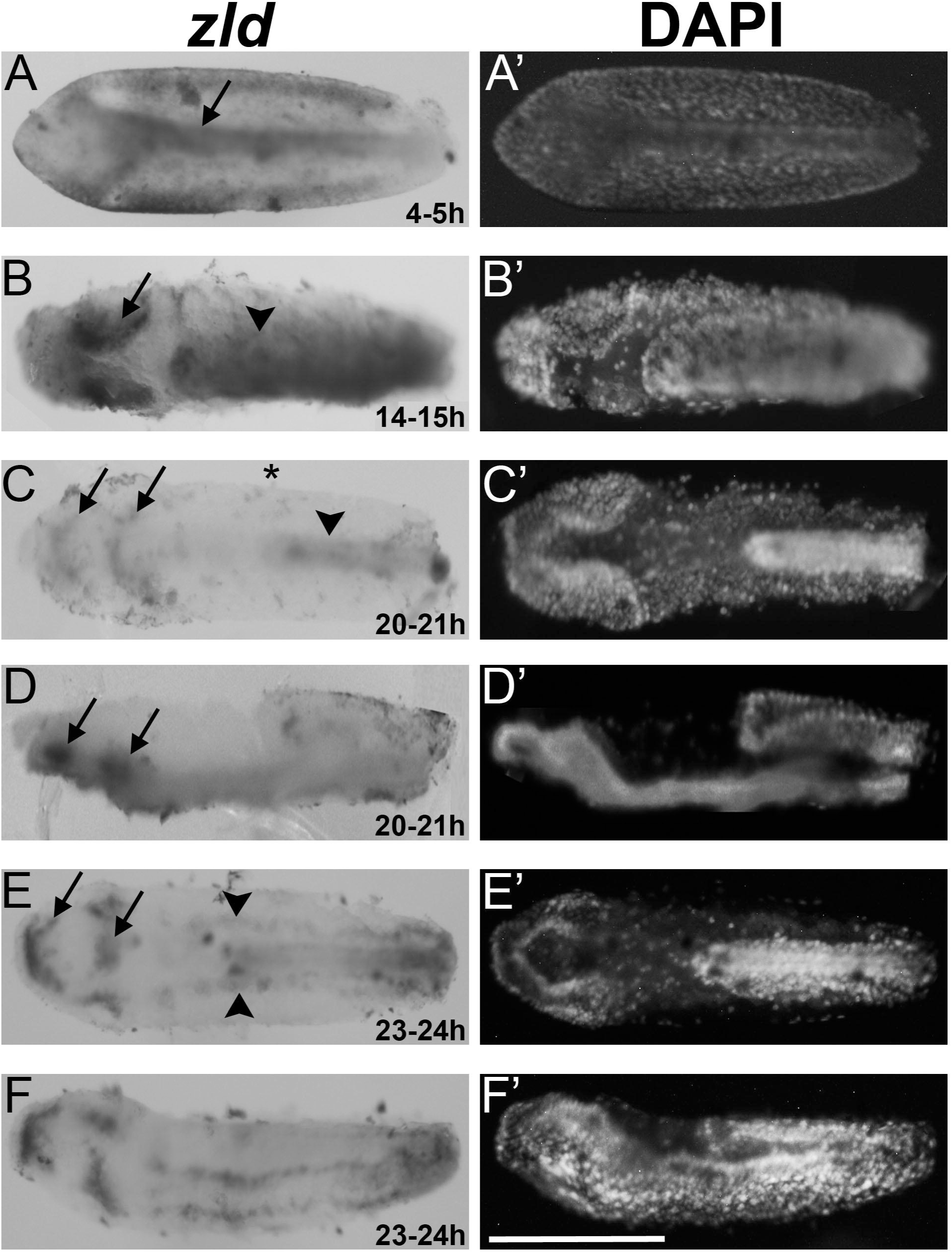
Expression pattern of the transcription factor *zelda* during *Aedes aegypti* embryonic development. (A, A’) Ventral view, embryo at 4–5 hours AEL. Embryo at 4–5 hours showing prominent staining in the prospective mesoderm (arrow) and in the lateral regions. (B, B’) Dorsal view of a 14–15-hour AEL embryo with strong staining in the head region (arrow) and germ band (arrowhead). (C, C’) Dorsal view of a 20–21-hour AEL embryo with staining in the anterior (head, arrows) and posterior (germ band, arrowhead) regions, as well as diffuse staining in the serosal layer (asterisk). (D, D’) Lateral view of the same 20–21-hour AEL embryo as in C, C’, highlighting head staining (arrows). (E, E’) Dorsal view of a 23–24-hour AEL embryo showing staining in the head (arrows) and along the middle region, specifically in parallel-aligned neuroblasts (arrowheads). (F, F’) Lateral view of the same 23–24-hour AEL embryo as in E, E’. Scale bar: 200 µm. Images captured using a Leica M205 stereomicroscope.

### RT-qPCR analysis of *mlpt, cact*, and *zld* expression during *Ae. aegypti* oogenesis and embryogenesis

The expression of the three developmentally regulated genes (*mlpt, cact*, and *zld*) was analyzed by RT-qPCR method during oogenesis and early embryogenesis. Ovariole growth in mosquitoes is triggered by blood feeding, and gene expression was examined in the ovary (Figure 5 A) and carcass (Figure 5 B) at 24, 48, and 72 hours post-blood feeding. Ovary (Figure 5 A) and carcass (Figure 5 B) expression profiles of the three genes showed a similar but not identical profile. Both *mlpt* and *cact* were expressed in the ovary and carcass at all three time points, with relative mRNA levels increasing significantly at 48 and 72 hours. Relative mRNA levels of both genes are higher at 48 and 72 hours than 24 hours after blood feeding. In contrast, *zld* expression was not detected in the same cDNA preparations, suggesting either very low mRNA levels or a lack of expression during oogenesis. RT-qPCR analysis during early embryogenesis revealed that *mlpt* and *cact* mRNAs are maternally deposited, as they were detected as early as 0–1 hour after egg laying (AEL; Figure 5 C). Expression of *cact* increased notably between 3–4 and 4–5 hours AEL, indicating the onset of zygotic transcription (Figure 5 C). Interestingly, *zld* mRNA was absent at 0–1, 1–2, and 2–3 hours AEL, confirming that it is not maternally deposited. Instead, *zld* expression was first detected between 3–4 hours AEL and increased sharply between 4–5 and 5–6 hours AEL, coinciding with cellularization and gastrulation. Overall, our RT-qPCR corroborates our *in situ* hybridization data (Figures 1-3) showing that the pTF *zld* is not maternally expressed and that the dorsoventral gene *cact* functions both maternally and zygotically during *Ae. aegypti* embryogenesis.

**Figure 5:**
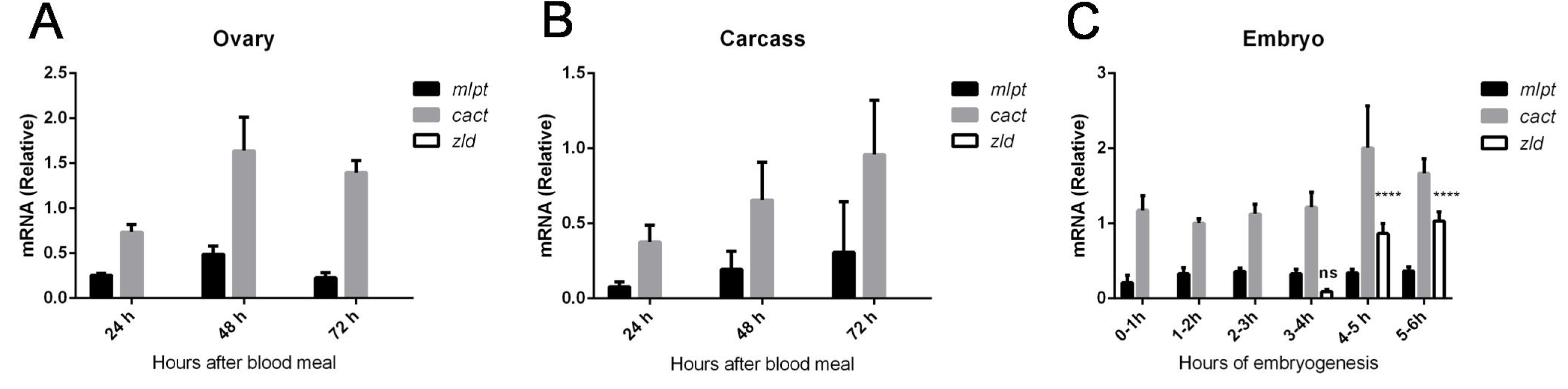
RT-qPCR analysis shows that *cact* and *mlpt* are maternally expressed, while *zld* expression is strictly zygotic. (A-C) Expression of *cact*, *mlpt* and *zld* during oogenesis and early embryogenesis (0-6 hours after egg lay). (A) Ovary (B) Carcass. (C) Expression in one-hour intervals, 0-1, 1-2, 2-3, 3-4, 4-5 and 5-6 hours. Experiments were performed in three biological replicates and error bars represent standard deviations. Graph data were analyzed using using the Kruskal-Wallis test. Differences were considered significant whenever p ≤ 0.05. ns: non-significant (p > 0.05); ****p ≤ 0.0001.

### Transcriptomic Analysis of *Ae. aegypti* Embryogenesis: Absence of Maternal *zld* Transcripts and Presence of Other Pioneer Transcription Factors

Previous studies have published transcriptomes for the earliest stages of *Ae. aegypti* embryogenesis, particularly spanning 0 to 7 hours of development (Biedler et al. 2012, Yoon et al. 2019). We compared the expression profiles of *zld, cact, mlpt*, and other transcription factors (TFs) implicated in genomic activation during the maternal-to-zygotic transition (MZT), including *Clamp, Trl-GAGA, opa, and grh*. Transcriptome analysis confirmed the absence of maternal *zld* transcripts (Figure 6, 0–3 hours) and the presence of zygotic *zld* transcripts beginning at 5–7 hours AEL. *cact* expression was observed throughout all developmental stages analyzed, confirming both maternal and zygotic contributions. Interestingly, *mlpt* displayed a similar profile to zld, with only zygotic transcripts detected.

**Figure 6:**
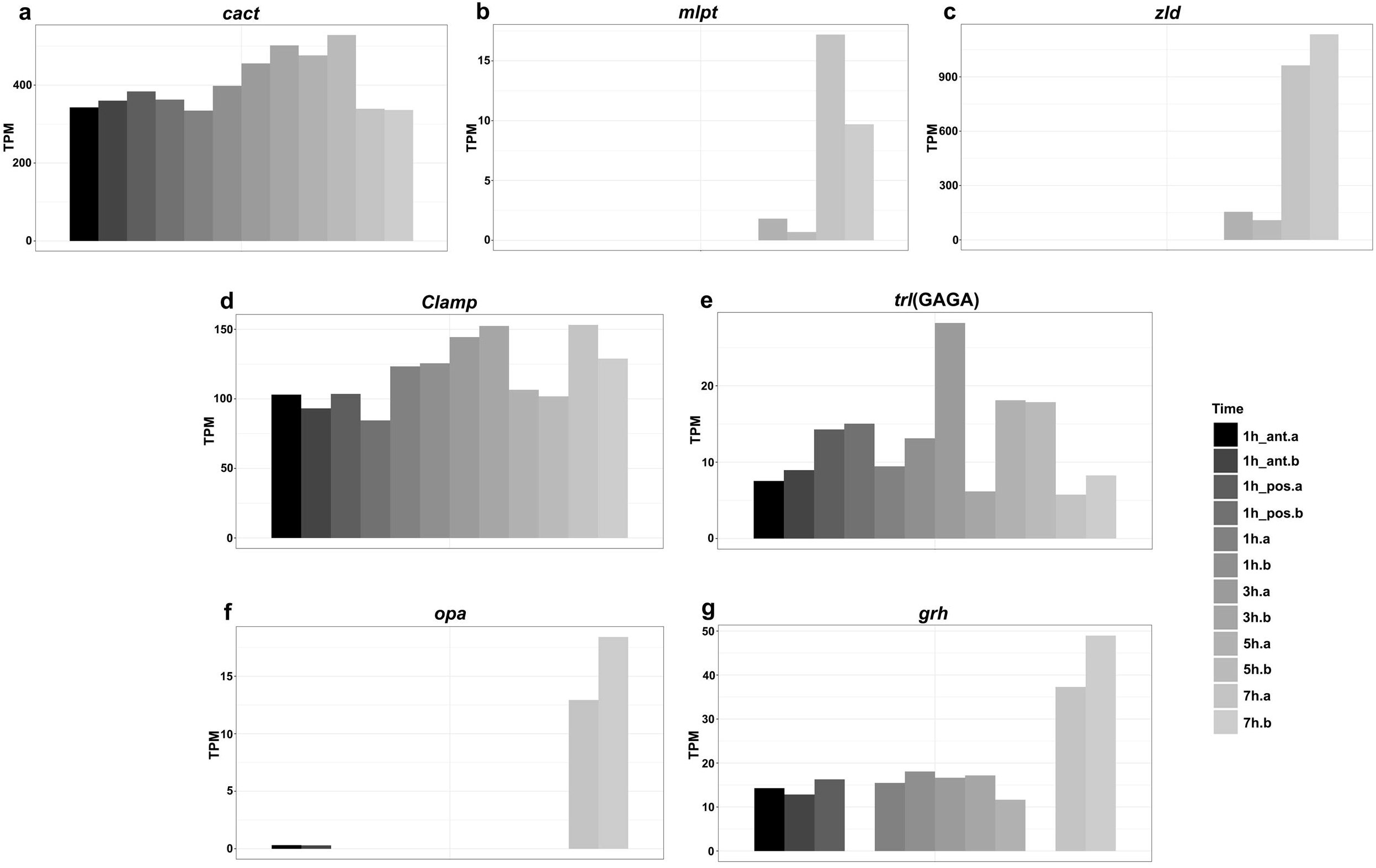
Expression profile of candidate genes responsible for genomic activation during *Ae. aegypti* maternal to zygotic transition and other developmental genes. a) *cactus (cact)*, b) *mlpt*, c) *zelda* (*zld*), d) *Clamp*, e) *Trl-GAGA*, f) *opa* and g) *grh*. 1-7h: hours of development of the entire body of the embryos, ant: anterior pole of freshly laid eggs, pos: posterior pole of freshly laid eggs and a-b: biological replicates. 0-1 hour (transcriptome data from Yoon et al. 2019), the remaining transcriptomes obtained from Bioproject PRJEB52866.

These transcriptome findings corroborate our *in situ* hybridization and RT-qPCR results, establishing that *zld* is strictly zygotically expressed in *Ae. aegypti*. Previously, Biedler et al. 2012 provided evidence that a domain remarkably similar to TAGteam was identified in *Ae. aegypti* genome, and initial data suggesting that the single ortholog of *zld* in *Aedes aegypti* was not maternally deposited. Thus our data show that *zld* does not function as the pioneer transcription factor (pTF) required for zygotic genome activation in this species. Given the reported roles of *Clamp, Trl-GAGA, opa* and *grh* as critical TFs for MZT in *Drosophila melanogaster* (Soluri et al. 2020, Duan et al. 2021, Gaskill et al. 2021), we analyzed their expression profiles in the *Ae. aegypti* transcriptomes. Our results reveal that *Clamp, Trl-GAGA,* and *grh* are expressed both maternally and zygotically, suggesting that one or more of these TFs may act as pTFs during mosquito zygotic genome activation.

Altogether, our study advances the understanding of early *Ae. aegypti* embryogenesis. First, we identified two distinct zygotic domains of *cact* expression in the mesoderm, indicating the presence of specialized mesodermal subtypes in mosquitoes. Second, *mlpt* is expressed in a broad gap domain during cellularization, resembling expression patterns in basally branching insect groups such as Coleoptera and Hemiptera, but differing from higher Diptera like fruit flies, where *mlpt* functions only at later embryonic stages. Third, the absence of maternal *zld* expression suggests that *zld* does not perform the same role in *Ae. aegypti* MZT as in *D. melanogaster*. By using a comparative multispecies approach (Nunes-da-Fonseca 2022), our findings contribute to the evolutionary understanding of MZT and early embryonic patterning, highlighting *Ae. aegypti* as a valuable model for studying these processes.

## Acknowledgements

The authors would like to thank the Integrated Unit of Microscopy and the Integrated Genomics Facility from the Instituto de Biodiversidade e Sustentabilidade (NUPEM/Federal University of Rio de Janeiro) for kindly providing access and support for the use of these facilities. We also are grateful to Juliana Beraldini for the help with animal care.

## Author Contributions

R.C.,D.S. and M.U. conducted research and investigation, writing and original draft preparation. L.R., D.S., J.J.M.M., M.A.A. conducted research and investigation; M.F.C.L. funding acquisition and research supervision and investigation; R.N.-d-F.: conceptualization ideas, conducting research and investigation, writing and original draft preparation, funding acquisition.

