## Supplemental Table 1 for "Analysis of gene expression in *Aedes aegypti* suggests changes in early genetic control of mosquito development"

**Supplementary Table 1- Primers used for the synthesis of antisense probes for *in situ* hybridization, double-stranded RNA synthesis and Real time PCR (RT-qPCR)**

| Primer name Primer sequence | | Product size | Application |
| --- | --- | --- | --- |
| *Aae*-*zld*-F1 | ggccgcggATCAACCGGTGAAGTCGTTC | 556 bp | *In situ* and dsRNA |
| *Aae*-*zld*-R1 | cccggggcGGAACCGTTGACGAAGTGTT |  |  |
| *Aae*-*mlpt*-F | ggccgcggCGAGAATTATGGCCTTGGAA | 527 bp | *In situ* and dsRNA |
| *Aae*-*mlpt*-R | cccggggcACGAACCGTCAACCAAAAAT |  |  |
| *Aae*-*cact*-F | ggccgcggATCAAAAGGAGGAGCGTCG | 706 bp | *In situ* and dsRNA |
| *Aae*-*cact*-R | cccggggcTGAACCCGCTGAGTCAAGAT |  |  |
| *Aae*-*zld*-QF1 | GGAAATCTAACGAAGCTAGAAGACC | 141 bp \| | RT-qPCR |
| *Aae*-*zld*-QR1 | GATCTGGTTGTCCACTTGATTTAAC |  |  |
| *Aae*-*mlpt*-QF1 | GATCTAACCAACATAGGAGCCTACA | 146 bp \| | RT-qPCR |
| *Aae*-*mlpt*- QR1 | AAGGAGTCCATCTTTGAGTCGTAGT |  |  |
| *Aae*-*cact*- QF1 | TCTTGCGTTGAAGTGAGTGG | 171 bp | RT-qPCR |
| *Aae*-*cact*- QR1 | GACCCTCTGAAAGGGAAAGG |  |  |
| *Aae-Rp49-QF1* | GCTATGACAAGCTTGCCCCCA | 189 bp \| | RT-qPCR |
| *Aae-Rp49-* QR1 | TCATCAGCACCTCCAGCTC |  |  |

Universal primers containing T7 promoter sequences at one end, in the case of probe templates for *in situ* hybridization, or T7 promoter sequences at both ends, in the case of dsRNA synthesis templates (Sequences: T7 universal primer 3 'agggatcctaatacgactcactatagggcccggggcg / T7 universal primer 5 'gagaattctaatacgactcactatagggccgcgg).
